# Building a literature knowledge base towards transparent biomedical AI

**DOI:** 10.1101/2024.09.22.614323

**Authors:** Yuanhao Huang, Zhaowei Han, Xin Luo, Xuteng Luo, Yijia Gao, Meiqi Zhao, Feitong Tang, Yiqun Wang, Jiyu Chen, Chengfan Li, Xinyu Lu, Tiancheng Jiao, Jiahao Qiu, Feiyang Deng, Lingxiao Guan, Haohong Shang, Fan Feng, Thi Hong Ha Vu, Thomas Bate, Dongxiang Xue, Jean-Philippe Cartailler, Michael Stitzel, Shuibing Chen, Marcela Brissova, Stephen Parker, Jie Liu

## Abstract

As artificial intelligence (AI) continues to advance and scale up in biomedical research, concerns about AI’s trustworthiness and transparency have grown. There is a critical need to systematically bring accurate and relevant biomedical knowledge into AI applications for transparency and provenance. Knowledge graphs have emerged as a powerful tool that integrates heterogeneous knowledge by explicitly describing biomedical knowledge as entities and relationships between entities. However, PubMed, the largest and most comprehensive repository of biomedical knowledge, exists primarily as unstructured text and is under utilized for advanced machine learning tasks. To address the challenge, we developed LiteralGraph, a computational framework to extract biomedical terms and relationships from PubMed literature into a unified knowledge graph. Using this framework, we established the Genomic Literature Knowledge Base (GLKB), which consolidates 14,634,427 biomedical relationships between 3,276,336 biomedical terms from over 33 million PubMed abstracts and nine well-established biomedical repositories. The database is coupled with RESTful APIs and a user-friendly web interface that makes it accessible to researchers for various usages. We demonstrated the broad utility of GLKB towards transparent AI in three distinct application scenarios. In the LLM grounding scenario, we developed a Retrieval Augmented Generation (RAG) agent to reduce LLM hallucination in biomedical question answering. In the hypothesis generation scenario, we elucidated the potential functions of RFX6 in type 2 diabetes (T2D) using the vast evidence from PubMed articles. In the machine learning scenario, we utilized GLKB to provide semantic knowledge in predictive tasks and scientific fact-checking.

## 1 Introduction

As artificial intelligence (AI) continues to advanced and scale up, its applications in biomedical research have quickly thrived [1, 2, 3]. However, concerns about AI’s trustworthiness and transparency have also grown [4], as knowledge embedded in the AI models is often difficult for humans to trace or verify [5]. Additionally, these models are prone to hallucinations and frequently lack knowledge provenance [6]. This is particularly problematic in knowledge-intensive applications such as biomedical question answering, experimental data interpretation, and hypothesis generation [7, 8]. Therefore, there is a critical need to systematically bring accurate and relevant biomedical knowledge into AI applications for transparency and provenance.

However, biomedical knowledge is loosely and heterogeneously manifested in tabular data, unstructured text, and images, making it difficult to consolidate into a uniform data structure. Knowledge graphs have emerged as a powerful tool that integrates heterogeneous knowledge by describing biomedical knowledge as entities and relationships between entities. Although various biomedical knowledge graphs have been developed to model functional relationships among genomic entities (e.g., genes, proteins, pathways, and GO terms) and biomedical terms (e.g., drugs and diseases) [9, 10, 11, 12, 13], PubMed, the largest and most comprehensive repository of biomedical knowledge, exists primarily as unstructured text.

To address the challenge, we developed a computational framework LiteralGraph to extract biomedical terms and relationships from PubMed literature into a unified knowledge graph. The computational framework standardizes the extracted knowledge by referencing controlled vocabularies from OBO ontologies and implements a well-defined BioCypher schema [14], ensuring compliance with FAIR principles. The yielded knowledge graph is called *Genomic Literature Knowledge Base* (GLKB), which consolidates 14,634,427 biomedical relationships between 3,276,336 biomedical terms from over 33 million PubMed abstracts and nine well-established biomedical repositories. The database is coupled with RESTful APIs and a user-friendly web interface (https://glkb.dcmb.med.umich.edu/) that makes it applicable to different downstream applications. To demonstrate the broad utility of GLKB towards transparent AI, we identify *three* distinct application scenarios. First, we show that GLKB effectively grounds large-language models (LLMs) and reduces LLM hallucination via retrieval augmented generation (RAG) [15]. We developed a question-answering (QA) agent to systematically translate natural language queries into graph queries over GLKB, and generate answers with references to PubMed articles. Second, GLKB helps scientists in hypothesis generation stemming from their experimental data. We demonstrated this application in a use case of exploring the RFX6 gene function in type 2 diabetes, in which we identified potential genegene and gene-disease associations using the vast evidence from PubMed literature [16]. Last but not least, GLKB allows machine learning (ML) models to effectively access literature through the semantic embeddings of the biomedical terms in GLKB. These embeddings are not only accurate in terms of representing the underlying biomedical concepts, but also improve ML tasks such as functional predictions and scientific fact-checking.

## 2 Results

### 2.1 GLKB connects diverse biomedical knowledge from PubMed articles with external biomedical repositories

In order to systematically extract biomedical knowledge from PubMed literature and make the extracted knowledge coherent with existing ontologies and repositories, we developed a computational framework LiteralGraph (Supplementary Fig. S1). Using LiteralGraph, we linked PubMed literature with a vast collection of biomedical repositories. The resulting knowledge graph, GLKB, particularly focuses on the genomic domain in which many well-defined ontologies are established (Fig. 1A, Supplementary File 1).

**Figure 1.**
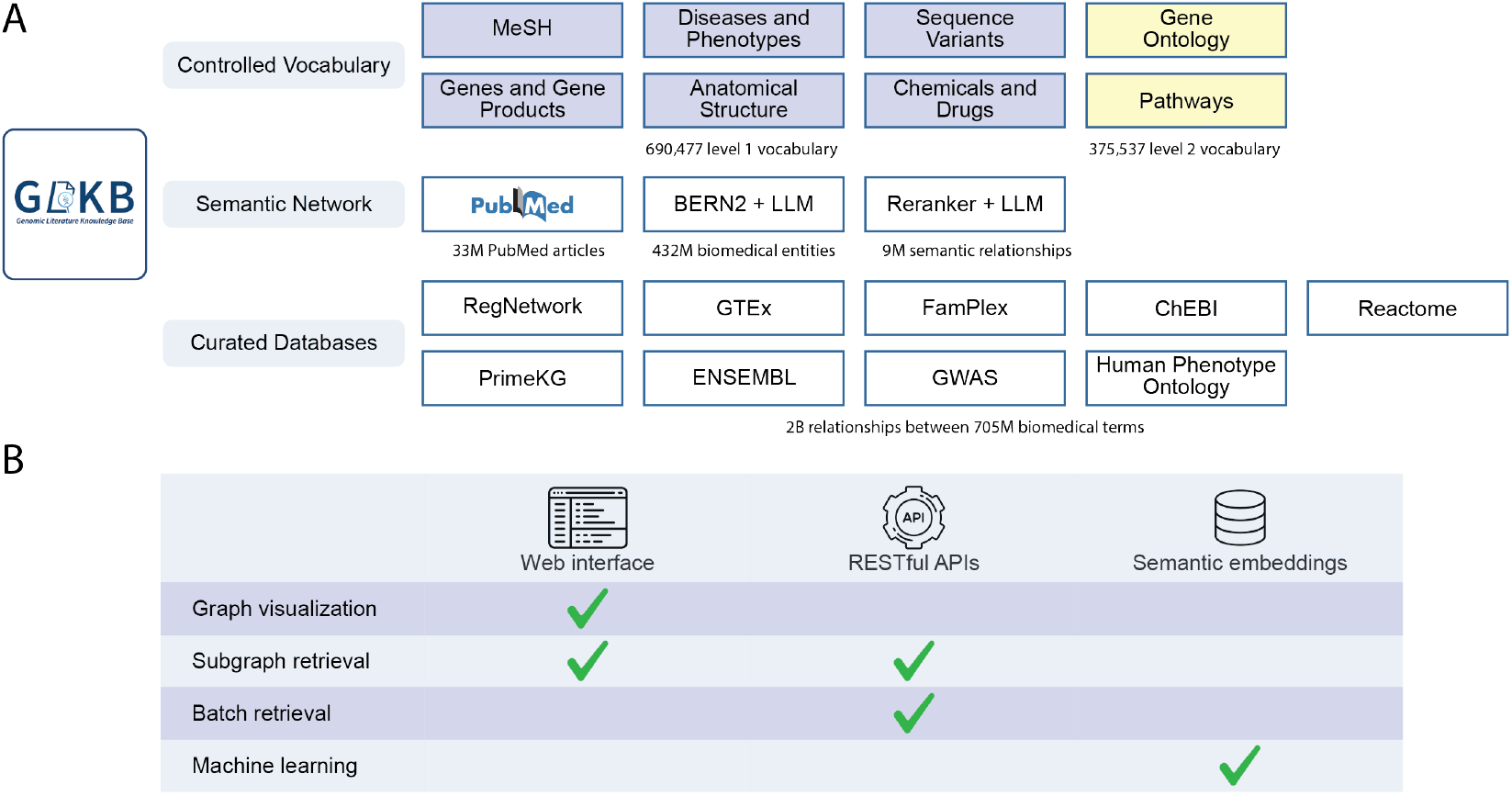
GLKB integrates a semantic network and curated databases using a controlled vocabulary. **A**. The scope of GLKB includes three parts, two levels of controlled vocabulary adapted from biomedical ontologies, a semantic network containing PubMed articles, biomedical entities, and semantic relationships, and a curated network that integrates biomedical databases annotated by domain experts. **B**. GLKB is accessible via three approaches, including the web interface, the APIs, and the data dump of the semantic embeddings.

The knowledge graph integrates two levels of controlled vocabulary. The first level includes foundational biomedical terms such as MeSH, genes and gene products, diseases and phenotypes, chemicals and drugs, sequence variants, and anatomical structures, all of which are directly mapped to entities extracted from the literature. The second level includes affiliated ontologies, which are linked specifically to certain types within the first-level vocabulary, including gene ontologies and pathways. In total, this controlled vocabulary encompasses 690,477 level-one terms and 41,311 level-two terms, with a comprehensive 375,537 ontology mappings identified.

Biomedical entity extraction was conducted using the BERN2 model [17], from which we extracted 432,417,310 biomedical entities across 33,403,054 PubMed articles. Of these, 341,319,268 entities (78.93%) were successfully mapped to at least one biomedical term in the controlled vocabulary. Following large language model (LLM) evaluation, 257,222,596 entities (75.36%) were validated and mapped to 500,272 distinct biomedical terms, while 19,660 terms were blacklisted due to inaccurate mappings. Among the 19,407,338 biomedical term pairs with at least five co-occurrences across PubMed articles, 9,286,411 pairs (47.85%) contained at least one article supporting a direct relationship. The relationships for these term pairs were then summarized from the most relevant articles using LLMs.

In addition to the extracted semantic knowledge graph, GLKB integrates nine human-curated databases, linking 56,930 biomedical terms with 3,795,248 structural and functional relationships. This integration provides a rich background of curated knowledge, offering a robust basis for further exploration and analysis. Because the curated network connects level-one and level-two vocabulary, we only connect level-one vocabulary to the articles to avoid redundancy.

GLKB is accessible in three ways, a user-friendly web interface, RESTful command-line APIs, and a data dump of the semantic embeddings (Fig. 1B). The web interface allows users to search and visualize GLKB’s graph structure. The APIs enable users with computational expertise to retrieve and analyze the PubMed literature at scale. The data dump allows a user to load semantic knowledge from PubMed to any machine learning model.

### 2.2 GLKB reduces LLM hallucination in biomedical domains

LLMs face two major challenges in biomedical research, the generation of fabricated results, often referred to as hallucinations [5, 6], and the need for frequent retraining to maintain currency with emerging knowledge [18]. GLKB addresses both issues by providing biomedical contexts from PubMed via retrieval-augmented generation (RAG) frameworks [19, 20]. To comprehensively support distinct applications, GLKB offers three approaches to retrieve biomedical knowledge in different needs (Fig. 2A). The first approach is the textual features of the entities, which enable precise retrieval of the biomedical concepts within the controlled vocabulary. The second approach is the semantic embeddings, which facilitate flexible retrieval of PubMed articles that contain specific research topics. The last approach is the graph structure, which supports retrieving complicated schematic associations among multiple entities. Using APIs, GLKB can be seamlessly integrated into RAG methods via all three approaches [21], enhancing LLM credibility and reliability.

**Figure 2.**
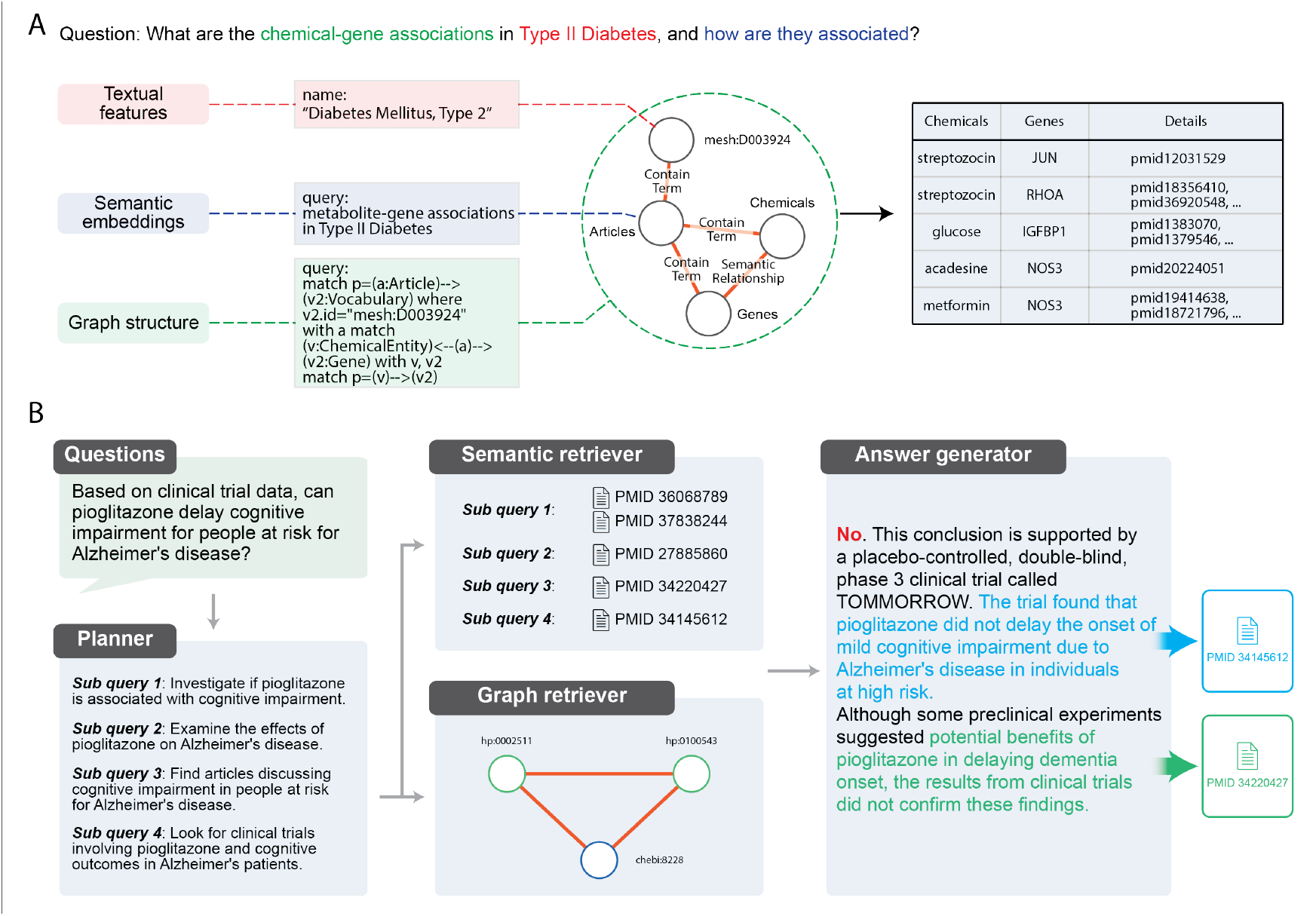
GLKB reduces LLM hallucination in biomedical question answering. **A**. Biomedical knowledge can be retrieved from GLKB via three approaches, namely textual features on entities, semantic embeddings, and graph structure. Combining knowledge in these formats enables GLKB to answer biomedical questions comprehensively. **B**.An GLKB QA agent is implemented to enhance LLM in biomedical question answering using PubMed articles and biomedical relationships. Utilizing domain knowledge retrieved from GLKB, the LLM agent is able to interpret the result with real-world biomedical evidence and pointers to two PubMed articles.

We tested GLKB in a biomedical QA task using GPT-3.5-turbo with and without the context from GLKB [22]. The results were evaluated by accuracy and *F*_1_ score on two biomedical question-answering datasets, PubMedQA and BioASQ, in a zero-shot setting [23, 24]. The two datasets contain 500 and 1,271 research questions, respectively, that can be answered with “yes”, “no”, or “maybe”. In addition to correct answers, PubMedQA also provides the original PubMed articles from which the questions were derived, making it suitable for evaluating GLKB’s retrieval performance. On the PubMedQA dataset, GLKB correctly retrieves the corresponding PubMed abstracts as the most relevant results for all 500 test questions using cosine similarities and significantly improves GPT-3.5-turbo’s performance with the retrieved context (Supplementary File 2). GLKB significantly improved performance, increasing accuracy from 0.62 to 0.76 and *F*_1_ score from 0.44 to 0.54 on PubMedQA, similar to human performance. On BioASQ, accuracy rose from 0.80 to 0.88 and *F*_1_ score from 0.70 to 0.78, confirming GLKB’s ability to retrieve essential biomedical domain knowledge and enhance LLMs in biomedical QA tasks (Supplementary File 3).

We further standardized the process in the GLKB QA agent which consists of four key components as follows (Fig. 2B). The *planner* decomposes input questions into discrete steps, addressing each with knowledge from PubMed articles or curated biomedical repositories. This stepwise breakdown ensures that even complex queries are handled effectively. The *semantic retriever* extracts detailed knowledge from PubMed articles using GLKB’s multi-level indexes. The *graph retriever* accesses relevant information from GLKB’s curated repositories, focusing on node and relationship properties to complement the semantic retriever by expanding the scope of knowledge retrieval. Finally, the *answer generator* synthesizes the retrieved information from multiple subqueries to generate well-grounded responses with proper references. Together, these components enable the GLKB QA agent (available via GLKB API) to deliver interpretable, reliable answers with references to PubMed articles and reduce LLM hallucinations.

### 2.3 GLKB facilitates biomedical hypothesis generation from PubMed evidence

Experimental scientists often struggle to interpret experimental data and generate hypotheses for new experiments, which requires an extensive review of the related literature. GLKB simplifies this process by systematically retrieving associations between biomedical terms. Here, we demonstrate how an experimental scientist can use GLKB to elucidate a gene’s function by leveraging extensive PubMed evidence. *RFX6* has recently been identified as a hub gene for beta cell function, but its relationship to other Type 2 diabetes (T2D) related genes is not well understood [16]. Using GLKB, scientists can identify T2D-related genes that are associated with *RFX6*, and further explore their functional groups.

The process involves two steps (Fig. 3A). First, we screened 2,675 differentially expressed genes (DEGs) from an *RFX6*-knockdown dataset [16] with the hope of identifying key genes modulated by *RFX6*. With GLKB, we identified ten significant genes based on strong literature evidence with a co-occurrence test (Method Section 4.6). Next, we analyzed how these genes are related to T2D by testing their co-occurrence with 7,018 DEGs from 20 T2D-related gene-KO cell lines using GLKB [25], identifying 68 T2D-related genes, seven of which overlapped between the two steps. We subsequently combined the two gene sets into 72 genes that potentially associated with both *RFX6* and T2D (Supplementary File 4). To further investigate the role of these identified genes in T2D, we selected 36 putative T2D causal genes, classified as “causal” based on the “curated approach probability”, from a total of 132 effector genes in the Type 2 Diabetes Knowledge Portal (T2DKP) [26]. In particular, more than half (*n* = 19*/*36) of the T2D causal genes were included in our final set, highlighting the effectiveness of GLKB in identifying relevant biomedical associations.

**Figure 3.**
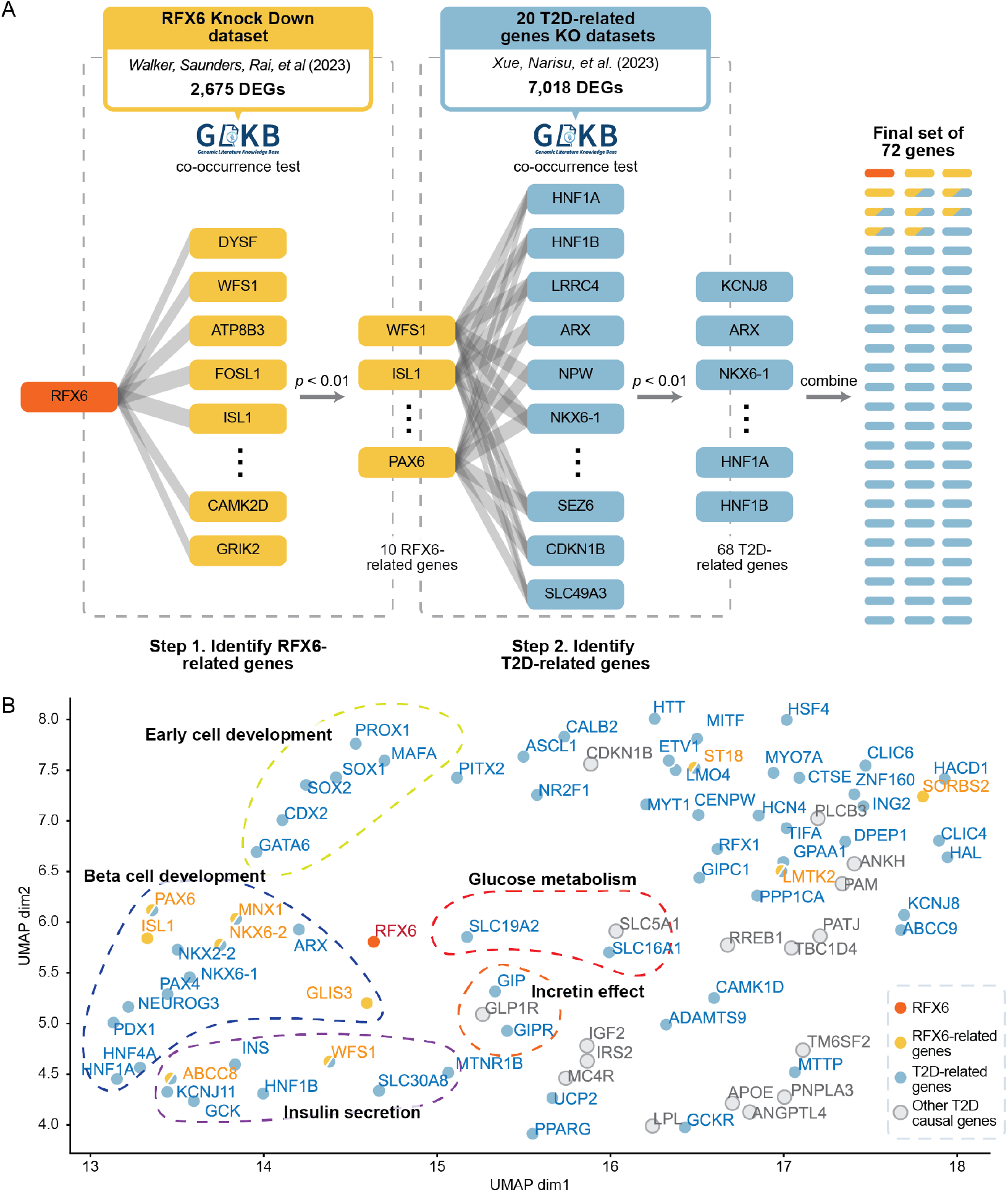
GLKB identified *RFX6*-related genes from PubMed articles and revealed their potential relationships to T2D. **A**. 10 *RFX6*-related genes (in yellow) were identified by GLKB from 2,675 DEGs in the *RFX6*-KD cell line, and 68 T2D-related genes were further found (in blue), including seven overlapped with the ten *RFX6*-related genes, associated with these *RFX6*-related genes from 7,018 DEGs in 20 KO cell lines where T2D-related genes were knocked out. In total, a combined final set of 72 genes related to the *RFX6*-T2D mechanisms was selected using GLKB. **B**. GLKB reflects the underlying mechanisms of the *RFX6*-related genes to T2D. The 72-gene final set is combined with the remaining 17 T2D causal genes from the Type 2 Diabetes Knowledge Portal (in grey), and visualized according to their co-occurrence in PubMed articles. Genes with different biological functions are located accordingly, and the distances between different genes indicate the strength of associations between them. According to the result, *RFX6* is associated with beta cell functions (e.g., insulin secretion and glucose metabolism) and beta cell development.

To further investigate the roles of these 72 *RFX6*-related genes in T2D, we analyzed their interactions with the remaining 17 *RFX6*-independent T2D causal genes from T2DKP as controls. The 89 genes were grouped based on co-occurrences in PubMed, revealing four functional groups (Fig. 3B). Genes with four types of functions were found in proximity to *RFX6*, highlighting different T2D mechanisms. The first group includes genes involved in beta cell development and differentiation, such as *PAX6, ISL1, MNX1*, and *NKX6-2*, along with T2D-related genes like *PDX1, HNF4A*, and *GLIS3* [27, 28]. *The second group involves genes like SLC19A2* and *SLC16A1*, key to glucose metabolism [29, 30, 31]. The third group includes incretin-related genes (*GIP, GIPR*, and *GLP1R*) that stimulate insulin secretion [32]. The fourth group includes genes such as *ABCC8, KCNJ11*, and *INS*, critical for insulin secretion and beta cell function, where mutations cause MODY [33, 34]. Another group, less related to *RFX6*, includes genes that regulate stem cell maintenance and early cell development, such as *GATA6* and *SOX2* [35, 36]. *Interestingly, most of the RFX6*-independent T2D causal genes (*n* = 15*/*17) were not closely associated with *RFX6*, suggesting it mainly influences insulin secretion and beta cell development. Although gene co-occurrence in literature does not always indicate similar biological functions, and only “early cell development” and “incretin effect” groups form statistically significant in DBSCAN clusters, these functional groups combined with human curation provide valuable insights for refining experimental hypotheses.

### 2.4 GLKB leverages PubMed semantic knowledge in machine learning tasks

Due to its unstructured nature, semantic knowledge in PubMed literature is difficult to retrieve and therefore underutilized in ML tasks. GLKB represents this knowledge as an explicit graph, allowing ML models to effectively access literature through embeddings. GLKB encodes three types of biological terms—genes, chemicals, and diseases—into 1,536-dimensional embeddings using its semantic network (Method Section 4.4). These semantic embeddings capture the functions of the corresponding biomedical terms (Fig. 4A). For example, gene embeddings cluster according to their GO terms and KEGG pathways (Fig. 4B). We validated the embeddings with two tasks.

**Figure 4.**
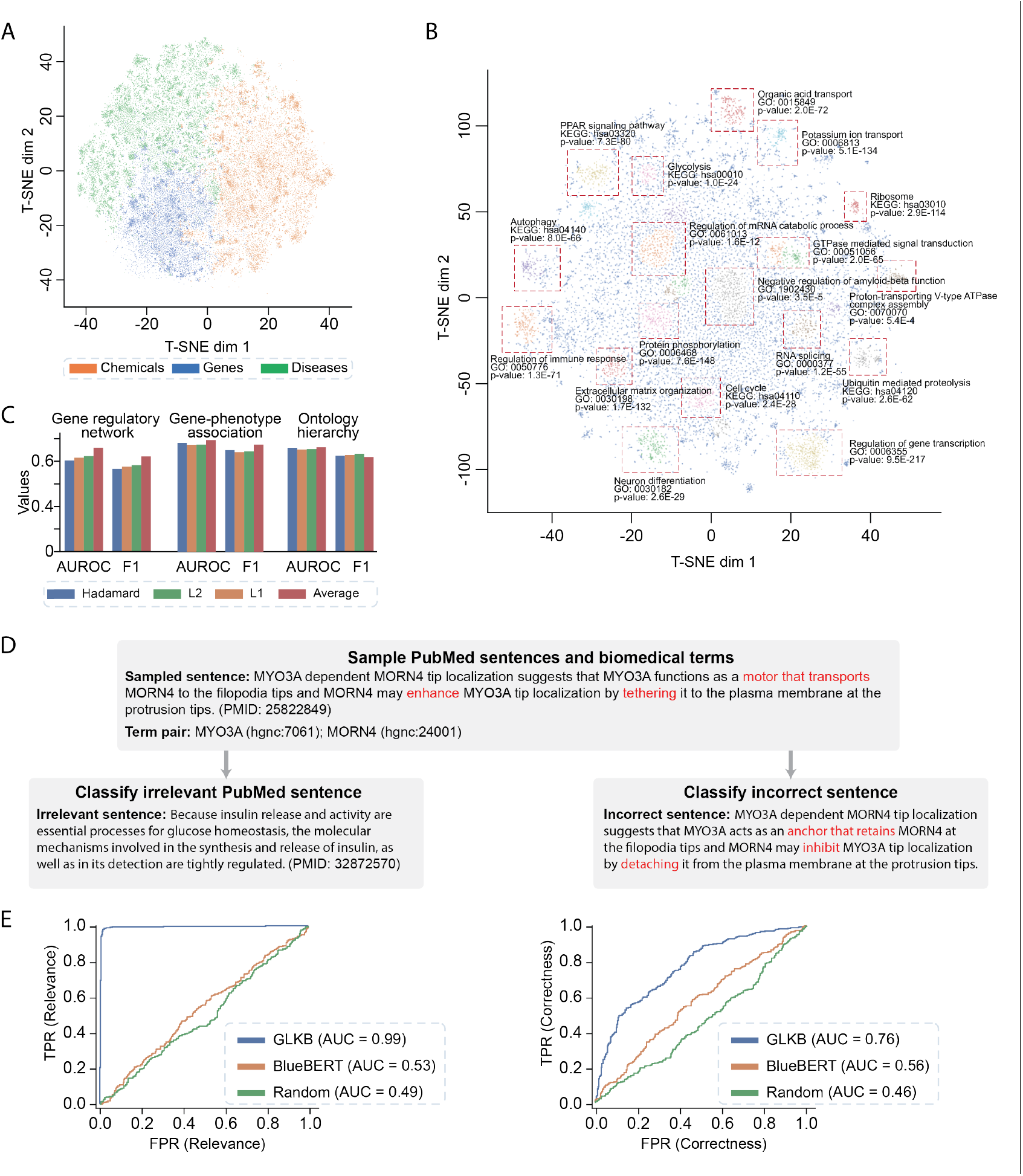
Semantic knowledge agrees with curated knowledge in GLKB. **A**. Semantic embeddings of genomic terms in GLKB visualized by T-SNE colored with genomic term types. **B**. Clusters in gene embeddings are highly associated with the biological functions of corresponding genes. Dot colors indicate gene clusters identified by DBSCAN. **C**. The semantic embeddings effectively predict curated biological relationships in genomic databases. The performance of predicting different types of human-curated knowledge using semantic embeddings is evaluated by AUROCs and *F*_1_ scores. Four pooling methods are used for prediction, including hardamard product, *L*_1_ norm, *L*_2_ norm, and average pooling. **D**. Examples of the scientific fact-checking task. In this experiment, we use the cosine similarity between biomedical term embeddings and different sentences to identify real relevant PubMed sentences against randomly sampled irrelevant sentences or fake sentences with incorrect knowledge. **E**. The semantic embeddings successfully distinguish real PubMed sentences against irrelevant sentences (left) and fake sentences (right), evaluated by AUROCs.

In the first task, we used the embedding to predict functional relationships between biomedical concepts. We trained logistic regression models to predict six types of relationships in the GLKB curated network, including gene regulatory networks, ligand-receptor pairs, and gene-disease associations (Fig. 4C, Supplementary File 5). The semantic embeddings well predicted all real-world relationships, suggesting the semantic embeddings reflect real-world biological functions.

In the second task, we used the embeddings for scientific fact-checking. Specifically, we used the semantic embedding to classify real 1,246 PubMed sentences from two negative sets: randomly selected PubMed sentences and fake sentences generated by GPT-3.5-turbo (Fig. 4D). The first set evaluates the semantic relevance of the embeddings, while the second set evaluates their scientific correctness. We expected higher cosine similarity between the embeddings and true sentences in both sets if they captured precise PubMed knowledge. The embeddings showed an AUROC of 0.99 in distinguishing real from irrelevant sentences, and 0.77 in differentiating real from fake sentences, outperforming BlueBERT (Fig. 4E) [37]. This confirms that the embeddings faithfully capture semantic relationships from PubMed.

## 3 Discussion

We developed a principled computational framework LiteralGraph which resolves existing ontologies, connects existing biomedical data repositories, extracts knowledge from PubMed, and produces a consolidated knowledge graph to capture the comprehensive knowledge explicitly. To our best knowledge, it is the first time that PubMed is systematically connected to existing data repositories and ontologies at the detailed knowledge level. Because knowledge is explicitly represented in a machine-learning ready format, these nodes, edges, and properties can be directly fed into computers for advanced, AI-powered downstream applications [38, 39]. For example, we applied GLKB to RAG agents for LLM grounding and utilized evidence from PubMed literature to facilitate the interpretation of experimental data for biomedical hypothesis generation. Additionally, we applied semantic embeddings to integrate PubMed semantic knowledge into various biomedical machine learning tasks, such as predicting biomedical associations between entities and scientific fact-checking. We expect that more AI applications can benefit from the use of the structural knowledge in GLKB.

When we developed LiteralGraph and GLKB, we aimed to meet the FAIR principles, namely findable, accessible, interoperable, and reusable, benefiting the research community in two ways [40]. First, we followed a well-defined schema based on BioCypher, which allows easy adaptation to different research domains and connecting with external data repositories and knowledge bases [14]. Second, the framework utilizes a modular architecture, which supports convenient data reusing and pipeline updating. The framework is adaptive and systematic in the sense that all input data and computational models are described via JSON files and the pipelines are shared with the community. When any component needs to be updated in the future, we only have to update the JSON files and rerun the corresponding modules. Therefore, we expect that LiteralGraph can be used to build knowledge graphs in other domains.

In this paper, we tested fundamental use cases of GLKB, which can be further extended. In the LLM integration task, GLKB, as both a PubMed vector store and a graphical database, seamlessly connects with LLM agents in multiple ways, such as generation-augmented retrieval (GAR) [41], hybrid retrieval [42], chain-of-thoughts reasoning [43], and tree-of-thoughts reasoning [44]. These strategies enhance LLM performance in more complex tasks such as clinical question answering, where the query typically involves clinical records and requires reasoning from multiple articles to generate the final answer. GLKB also provides a feasible platform for advanced hypothesis generation frameworks. For instance, instead of merely identifying significant biomedical term pairs, machine learning agents such as reinforcement learning (RL) models can be employed to generate more comprehensive hypotheses based on multi-hop trajectories in GLKB [45]. By integrating more advanced machine learning models with GLKB, we believe that GLKB can become an even more valuable resource for these models to systematically utilize the knowledge within PubMed.

## 4 Online methods

### 4.1 LiteralGraph framework

To integrate PubMed with existing biomedical repositories and knowledge graphs, we developed the *LiteralGraph* framework. This framework was designed to bridge the extensive PubMed literature with curated biomedical resources in a scalable, maintainable, and accurate manner.

The LiteralGraph framework operates in *four* steps (Fig. S1). First, ontologies are merged to construct a controlled vocabulary that defines the scope and serves as the backbone of the knowledge graph. This vocabulary establishes a standardized set of biomedical terms that ensures consistency and interoperability with other resources. Second, a local named entity recognition (NER) model is applied to PubMed abstracts to identify and map biomedical entities to the controlled vocabulary, creating a structured representation of the literature. Third, LLMs evaluate these mappings to filter out potential misclassifications. Specifically, LLMs evaluate 2,000 sampled entity-sentence pairs mapped to each biomedical term in the controlled vocabulary, flagging mappings that show a high proportion of mismatches. Finally, relationships are summarized between pairs of biomedical terms that co-occur in PubMed articles. For each term pair, the top 1,000 co-occurrence articles are ranked by a reranker model to assess relevance, and the LLMs then summarize these articles to describe the relationship between the terms.

### 4.2 Data sources of GLKB

#### Data retrieval

The articles in GLKB are obtained in XML format from the PubMed database. From each article, we extract various information such as the PubMed ID, title, abstract, keywords, authors, publication date, ISSNs of the published journal, citations, and publication types. Ontologies in GLKB are retrieved from NCBO BioPortal in CSV format, and for each genomic term, we extract its name, synonyms, Uniform Resource Identifier (URI), source ontology, MeSH tree location, cross-references, and NCBO LOOM mappings. Journals in GLKB are retrieved from the Medline database in TXT format, and we extract details such as JrID, NLM ID, title, NLM abbreviation, ISO abbreviation, online ISSNs, and printed ISSNs. Genomic databases in GLKB are obtained from GenomicKB, the respective data portals, or web servers in various formats [13]. We implemented a Python pipeline to automatically convert the raw tabular data to CSV format, which can be easily loaded into the Neo4j data loader.

#### Data preprocessing

To prepare the PubMed abstracts for entity and relation extraction, we performed two preprocessing steps: sentence splitting and coreference resolution. First, we identified sentences in the original abstracts using the NLTK sentence splitter [46]. Next, we applied the NeuralCoref 4.0 model in spaCy [47, 48] to identify and resolve coreferences in the separated sentences. Coreferences are tokens that refer to other tokens or sets of tokens (e.g., “they”, “it”, and “their”) in the same or another sentence.

#### Vocabulary construction

To construct the controlled vocabulary in GLKB, biomedical ontologies are first downloaded from the OBO foundry using the Python package PyOBO 0.10.6 [49]. We resolved redundant gene, chemical, and disease terms in the vocabulary. For genes, we used the NCBI gene dataset as a reference, revealing 28,493 general gene matches, 14,406 pseudogene matches, and 767 other matches from a total of 43,794 gene entries, with only 128 entries requiring manual alignment. The alignment of the NCBI gene dataset with the MeshTerm dataset revealed 117 gene matches and 3 pseudogene matches, indicating some contamination within the MeshTerm dataset. These multi-referencing issues primarily arose from synonyms or outdated gene symbols, with mismatched terms including ontology terms which should be relocated to an ontology vocabulary. For chemicals, a direct alignment in the MeshTerm subset successfully identified 7,846 duplicated entries. For diseases, all entries were matched with the UMLS system using the provided [50], identifying 21,694 duplicated entries, which were remapped to properly represent the connected concept. This initiative ensures that gene, chemical, and disease terms in GLKB are accurately linked and updated, enhancing its utility and reliability for bioinformatics research.

### 4.3 Information extraction methods

#### Named entity recognition

Biomedical entity extraction was conducted using the BERN2 model [17]. We utilized the Gilda normalizer to map the extracted entities to the GLKB’s controlled vocabulary [51]. To ensure correct entity mapping, we set up a constraint that the entity and the genomic term it mapped to must have the same label. For example, an entity labeled “ER” would be mapped to the term “endoplasmic reticulum” instead of “erdosteine” if it was recognized as an anatomical structure by the entity extractor. The mappings of 95,219 high-frequency biomedical terms that mapped to ≥30 articles were further screened. For each high-frequency biomedical term, 100 corresponding entities and sentences were sampled, and the mappings were evaluated using GPT-4 LLM. All mappings to a biomedical term were deleted if more than 5 samples were classified as incorrect or uncertain. As a result, 20,049 terms were deleted at this stage.

#### Relationship extraction

Relationships between biomedical term pairs were summarized from their cooccurring PubMed sentences. Specifically, these sentences were reranked using a BGE reranker model [52]. Relationships were then summarized from at most 100 reranked sentences with normalized score ≥0.6 and mapped to one of the eight relationship types from the BioRED corpus using GPT-4, including positive and negative correlations, associations, bindings, drug interactions, cotreatments, comparisons, and conversions[53].

#### Ontology mapping

To identify and reduce redundant biomedical terms across ontologies, 375,537 ontology mappings were generated in GLKB’s controlled vocabulary using AML [54] and Logmap [55].

### 4.4 GLKB embedding

#### Biomedical term embedding

We utilized Heterogeneous Graph Transformer (HGT) to generate 1,536-dimensional embeddings of biomedical terms [56]. Due to computational constraints, we were compelled to produce embeddings only for 87,624 biomedical terms including genes, chemicals, and diseases. Firstly, we generated the input feature from the top 20 abstracts and the definition of each biomedical term using the OpenAI text-embedding-3-small model. The semantic network containing the 75,170 terms was then fed into a two-layer HGT model and trained on a link prediction task that predicts the co-occurrence of these terms. The final biomedical term embeddings are generated by the trained model.

#### Abstract and sentence embedding

Semantic embeddings of 33,403,054 PubMed abstracts and 108,536,203 sentences were generated using the gte-large-en-v1.5 model [57]. Due to computational constraints, a subset of sentences was selected for embedding. Specifically, we trained a spaCy sentence classification pipeline on the PubMed 200K RCT dataset, selecting only sentences labeled as “Background”, “Results” or “Conclusion” for embedding [47, 58].

### 4.5 System design

#### Software systems

GLKB was designed as a property graph that supports heterogeneous nodes, including articles, entities, and genomic terms, as well as edges such as citation, hierarchical relations, and genegene regulations, each with properties. To implement the database, we chose the *Neo4j* graph management system [59], which allows us to efficiently implement the property graph model directly down to the storage level. Additionally, the system offers an efficient query language, Cypher [60], which supports constant-time traversals for both depth and breadth queries over data graphs. With these features, GLKB provides a powerful tool for querying and analyzing complex genomic relationships.

#### Database update

GLKB tracks the latest releases of all input data resources and updates accordingly every twelve months to coincide with the PubMed annual update (current version release 2023).

#### Web interface

The GLKB database is hosted on a web server at the Michigan Academic Computing Center (MACC) at the University of Michigan, providing web-based access to users. The system backend is implemented using the Flask web framework and the socket library, with the graph database powered by Neo4j. To communicate with the Neo4j database, the server program uses the Neo4j Python driver. For interactive network visualization of the graph database, we used the *Cytoscape* framework in the frontend [61], while the user interface is built using the *React* framework. The NGINX web server acts as the reverse proxy, routing internet requests to our server, and no user cookies are collected. To support concurrent access from multiple users, the GLKB web server is designed with multi-threading and subjected to stress testing to ensure reliable service.

### 4.6 Pearson’s chi-square test for biomedical term co-occurrence

Pearson’s chi-square tests are performed to identify significant associations between biomedical terms. Specifically, for two given terms *T*_*i*_, *T*_*j*_, their expected co-occurrence frequency is

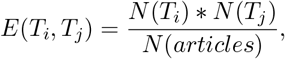

where *N* (*T*_*i*_) and *N* (*T*_*j*_) are the relevant articles of *T*_*i*_ and *T*_*j*_ respectively, and *N* (*articles*) is the total number of articles in GLKB. The test statistic is then calculated from the co-occurrence contingency table of the two terms. Clusters of the biomedical term co-occurrence are calculated by agglomerative clustering in python.

